# Bases of antisense IncRNA-associated regulation of gene expression in fission yeast

**DOI:** 10.1101/220707

**Authors:** Maxime Wery, Camille Gautier, Marc Descrimes, Mayuko Yoda, Valérie Migeot, Damien Hermand, Antonin Morillon

## Abstract

Antisense (as)lncRNAs can regulate gene expression but the underlying mechanisms and the different cofactors involved remain unclear. Using Native Elongating Transcript sequencing, here we show that stabilization of antisense Exo2-sensitivite lncRNAs (XUTs) results in the attenuation, at the nascent transcription level, of a subset of highly expressed genes displaying prominent promoter-proximal nucleosome depletion and histone acetylation. Mechanistic investigations on the catalase gene *ctt1* revealed that its induction following oxidative stress is impaired in Exo2-deficient cells, correlating with the accumulation of an asXUT. Interestingly, expression of this asXUT was also activated in wild-type cells upon oxidative stress, concomitant to *ctt1* induction, indicating a potential attenuation feedback. This attenuation correlates with asXUT abundance, it is transcriptional, characterized by low RNAPII-ser5 phosphorylation, and it requires an histone deacetylase activity and the conserved Set2 histone methyltransferase. Finally, we identified Dicer as another RNA processing factor acting on *ctt1* induction, but independently of Exo2. We propose that asXUTs could modulate the expression of their paired-sense genes when it exceeds a critical threshold, using a conserved mechanism independent of RNAi.

**AUTHOR SUMMARY:** Examples of regulatory antisense (as)lncRNAs acting on gene expression have been reported in multiple model organisms. However, despite their regulatory importance, aslncRNAs have been poorly studied, and the molecular bases for aslncRNAs-mediated regulation remain incomplete. One reason for the lack of global information on aslncRNAs appears to be their low cellular abundance. Indeed, our previous studies in budding and fission yeasts revealed that aslncRNAs are actively degraded by the Xrn1/Exo2-dependent cytoplasmic 5′-3′ RNA decay pathway. Using a combination of single-gene and genome-wide analyses in fission yeast, here we report that the stabilization of a set of Exo2-sensitive aslncRNAs correlates with attenuation of paired-sense genes transcription. Our work provides fundamental insights into the mechanism by which aslncRNAs could regulate gene expression. It also highlights for the first time that the level of sense gene transcription and the presence of specific chromatin features could define the potential of aslncRNA-mediated attenuation, raising the idea that aslncRNAs only attenuate those genes with expression levels above a “*regulatory threshold*”. This opens novel perspectives regarding what the potential determinants of aslncRNA-dependent regulation, as previous models in budding yeast rather proposed that aslncRNA-mediated repression is restricted to lowly expressed genes.

## INTRODUCTION

Eukaryotic genomes are pervasively transcribed [1], generating plenty of non-coding (nc) transcripts, distinct from the housekeeping rRNAs, tRNAs and sn(o)RNAs, and that are arbitrarily classified into small (< 200 nt) and long (≥ 200 nt) ncRNAs [2,3].

Long (l)ncRNAs are produced by RNA polymerase II (RNAPII), capped and polyadenylated, yet lack protein-coding potential [4,5], although this last point is subject to exceptions [6].

Several lines of evidence suggest that they are functionally important. First, IncRNAs show tissue-specific expression [7] and respond to diverse stimuli, such as oxidative stress [8], suggesting that their expression is precisely controlled. Second, several lncRNAs are misregulated in diseases including cancer and neurological disorders [9,10,11]. Furthermore, there is a growing repertoire of cellular processes in which lncRNAs play important roles, including X-chromosome inactivation, imprinting, maintenance of pluripotency and transcriptional regulation [12,13].

Several classes of lncRNAs have been described [2]. Among them, large intervening non-coding (linc)RNAs, which result from transcription of intergenic regions, have attracted a lot of attention as being involved in *cis*- and *trans*-regulation, mostly at the chromatin level, of genes important for development and cancer [13].

Another class of lncRNAs consists of antisense transcripts, that are produced from DNA strand antisense to genes [14]. Several examples of regulatory antisense (as)lncRNAs acting on sense gene expression in *cis* or in *trans* have been described in the budding yeast *Saccharomyces cerevisiae* [15,16,17,18,19,20,21], in the fission yeast *Schizosaccharomyces pombe* [22,23,24], in plant [25] and in mammalian cells [26,27].

Our previous studies in budding and fission yeasts revealed that aslncRNAs are globally unstable and are mainly targeted by the cytoplasmic 5′-3′ RNA decay pathway dependent on the Xrn1 and Exo2 exoribonucleases in *S. cerevisiae* [28,29] and *S. pombe* [30], respectively. Inactivation of Xrn1/Exo2 leads to the stabilization of a family of lncRNAs, referred to as Xrn1-sensititve Unstable Transcripts (XUTs), the majority of which are antisense to protein-coding genes [28,29,30].

Interestingly, in *S. cerevisiae*, we defined among these antisense (as)XUTs a subgroup for which the sense-paired genes (referred to as class 1) undergo antisense-mediated transcriptional silencing [28]. However, the molecular mechanism by which asXUTs could regulate sense gene expression remains largely unknown to date, still interrogating whether sense transcription is impaired at the initiation and/or elongation and/or termination stages, whether any post-transcriptional event is in play, and whether the epigenetic landscape contributes in the regulatory determinants. In addition, such a transcriptional aslncRNA-mediated regulation has not been documented yet in *S. pombe*.

Here we used Native Elongating Transcript sequencing (NET-Seq) to identify genome-wide in fission yeast the genes attenuated at the nascent transcription level upon stabilization of their paired-asXUTs. This so-called class 1 corresponds to highly transcribed genes, displaying marks of active transcription at the chromatin level. Mechanistic investigation on a model class 1 representative, the inducible catalase-coding gene *ctt1*, confirmed that it is transcriptionally attenuated upon oxidative stress when its paired-asXUT is stabilized, and the level of the attenuation correlates with the abundance of the asXUT. The attenuation is characterized by low RNAPII Ser5-phosphorylation (Ser5-P) and requires histone deacetylase (HDAC) activity and the conserved Set2 histone methyltransferase (HMT). Finally, we identified Dicer as an additional regulator of *ctt1* induction, acting independently of Exo2 and the asXUT. Together, our data support a model where asXUTs could modulate the expression of the paired-sense genes when it exceeds a critical threshold, using a conserved mechanism independent of RNAi.

## RESULTS

### Genome-wide identification of class 1 genes in fission yeast

In budding yeast, stabilization of asXUTs results into the attenuation, at the transcriptional level, of a subset of paired-sense genes, which are referred to as class 1 [28]. We recently annotated XUTs in fission yeast [30]. We observed that asXUTs accumulation correlates with down-regulation of the paired-sense mRNAs, at the RNA level [30], suggesting that the regulatory potential of asXUTs has been conserved across the yeast clade. However, whether this regulation occurs at the level of transcription in fission yeast remains unclear.

To define class 1 in *S. pombe*, we performed NET-Seq in WT and *exo2Δ* cells. Although global mRNA synthesis was found to be unchanged upon *exo2* inactivation (S1A Fig), differential expression analysis discriminated genes for which transcription in *exo2Δ* was significantly reduced (classes 1 & 2, n=723) or not (classes 3 & 4, n=4405). Within each category, we distinguished genes with (classes 1 & 3) or without (classes 2 & 4) asXUTs (Figs 1A-B; lists in Sl-4 Tables).

**Fig 1.**
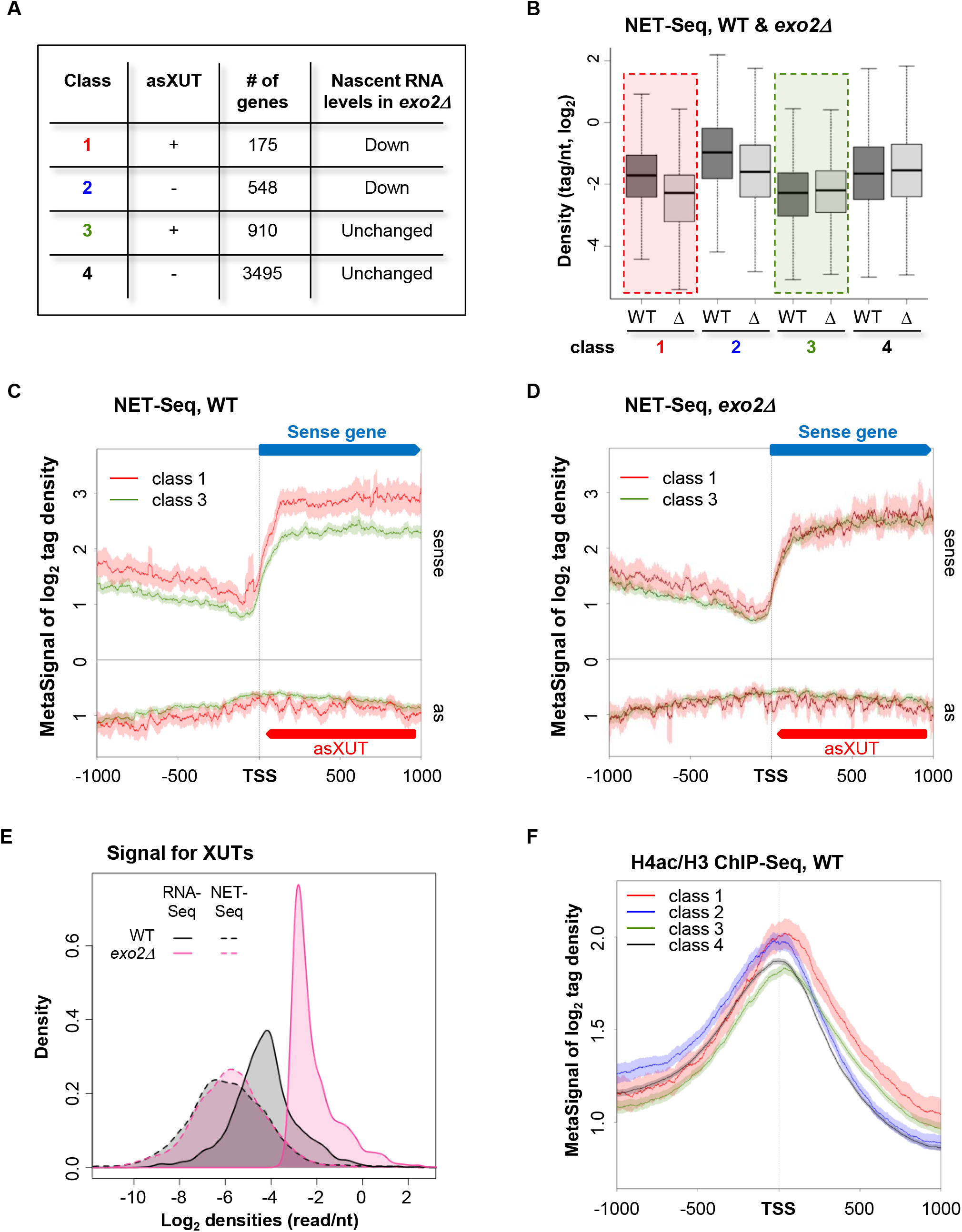
Genome-wide identification of class 1 genes in *S. pombe*. **A.** Transcriptional attenuation in *exo2Δ* cells. NET-Seq analysis was performed from biological duplicates of WT and *exo2Δ* cells. Data for the WT strain were previously described [30]. After sequencing, differential analysis discriminated genes showing significant (*P*<0.05) reduction of transcription (classes 1-2) or not (classes 3-4). Among them, classes 1 and 3 have asXUT. The number of genes for each class is indicated. **B.** Box-plot of nascent transcription (NET-Seq) signal for class 1-4 genes in WT (dark grey boxes) and *exo2Δ*(*Δ*; light grey boxes). **C.** Metagene view of NET-Seq signals along class 1 and 3 genes in WT cells. For each class, normalized coverage (tag/nt, log_2_) along mRNA transcription start site (TSS) +/− 1000 nt (sense) and the antisense (as) strand were piled up, in a strand-specific manner. Average signal for the sense and antisense strands was plotted for class 1 (red) and class 3 (green). The shading surrounding each line denotes the 95% confidence interval. **D.** Same as above in *exo2Δ*. **E.** Density-plot showing the global NET-Seq (dashed lines) and total RNA-Seq (solid lines) signals for XUTs in the WT (black) and *exo2Δ* (pink) strains. Total RNA-Seq data were previously described [30]. **F.** Metagene view of H4K5/8/12/16 acetylation (H4ac) for class 1 (red), class 2 (blue), class 3 (green) and class 4 (black) genes in WT cells. ChIP-Seq libraries construction and sequencing were previously described [30]. Metagene representation of signal for each class of genes was performed as above, in a strand-unspecific manner, using ratio of coverage (log_2_) for H4ac and H3. The shading surrounding each line denotes the 95% confidence interval.

Among the 723 genes transcriptionally attenuated in *exo2Δ*, 175 have asXUTs (class 1). Despite the proportion of class 1 genes among the attenuated genes is limited (24.2%), it is significantly higher than expected if presence of asXUT and sense gene attenuation were independent (Chi-square test, *P* = 0.03), suggesting that the attenuation could depend on the stabilized asXUTs, at least in some cases. On the other hand, the transcriptional down-regulation of class 2 (no asXUT) is likely to be an indirect effect reflecting the slow growth phenotype of the *exo2Δ* mutant [31]. Consistently, class 2 is significantly enriched for GO terms “ribosome biogenesis” (*P*=1.36e^−08^) and “cellular component biogenesis (*P*= 1.04e^−02^), and it is known that the expression of genes involved in these biological processes directly depends on the growth rate [32], Altogether, these observations suggest that for a subgroup of genes, stabilization of the asXUT might contribute to attenuate transcription of the paired-sense gene.

Both classes 1 and 3 have asXUTs, but only class 1 is transcriptionally attenuated upon asXUTs stabilization. This suggests the existence of specificities discriminating the two classes.

Indeed, in WT cells, class 1 is transcribed to higher levels than class 3 (Fig 1C), the latter actually showing the lowest transcription levels among the four classes (S1B Fig). In *exo2Δ*, transcription of class 1 falls to the low, basal level of class 3 (Fig 1D). Notably, transcription of XUTs antisense to class 1 and 3 genes is globally unaffected in the *exo2Δ* mutant (S1C Fig), indicating that XUTs accumulation in this context is due to the inactivation of their decay and not to a global increase of their synthesis (Fig 1E).

We also noted that in WT cells, the nascent antisense transcription signal surrounding the TSS of class 1 genes is higher than for class 3 (S1D Fig), suggesting that sense TSS overlap could constitute a factor for the potential regulatory activity of the XUTs antisense to class 1 genes.

At the chromatin level, class 1 shows a more pronounced nucleosome depletion in the TSS-proximal region than class 3 (S1E Fig), higher H3K14 (S1F Fig) and H4K5/8/12/16 acetylation (Fig 1F). When compared to the four classes, the levels of histone acetylation at the TSS-proximal region are similar for classes 1 and 2 (Fig 1F, see also S1F Fig).

Together, these results show that transcriptional attenuation correlates with asXUT stabilization in fission yeast, suggesting that asXUTs might be involved in the modulation of sense genes expression.

### Exo2-deficient cells are defective for *ctt1* induction upon oxidative stress

To further investigate the possibility that asXUTs can regulate expression of their paired-sense gene and to get insights into the underlying molecular mechanism, we characterized a class 1 gene, *ctt1*, and its paired-antisense *XUT0794* (Fig 2A, see also S2A Fig).

**Fig 2.**
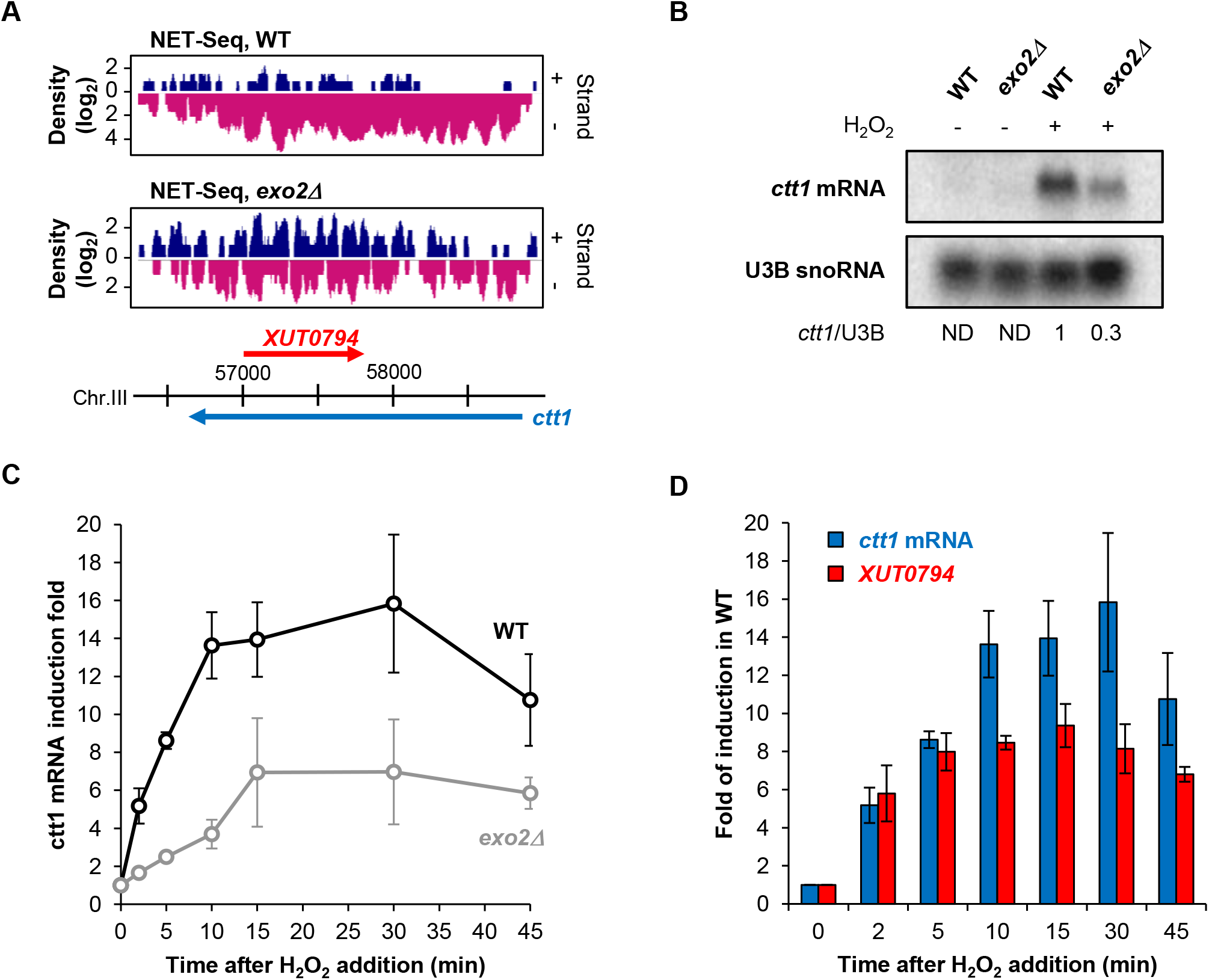
*ctt1* induction upon oxidative stress is impaired Exo2-deficient cells. **A.** Snapshot of nascent transcription (NET-Seq) signals along the *ctt1* gene in WT (upper panels) and *exo2Δ* (lower panels) cells. In each panel, the signal corresponding to the sense (+) and antisense (−) strand is shown in blue and pink, respectively. Blue and red arrows represent the *ctt1* mRNA and antisense *XUT0794*, respectively. NET-Seq data for the WT strain were previously described [30]. The snapshot was produced using VING [59]. **B.** Northern blot analysis of *ctt1* mRNA induction in WT and *exo2Δ* cells. YAM2400 (WT) and YAM2402 (*exo2Δ*) cells were grown to mid-log phase in rich medium and collected before or after addition of 1 mM H_2_O_2_ for 15 min. *ctt1* mRNA and U3B snoRNA were detected from total RNA using ^32^P-labelled oligonucleotides. Numbers represent the *ctt1* mRNA/U3B ratio (ND: not determined). **C.** YAM2400 (WT) and YAM2402 (*exo2Δ*) cells grown in rich YES medium to mid-log phase were collected 0, 2, 5, 10, 15, 30 and 45 minutes after addition of 1 mM H_2_O_2_. *ctt1* mRNA was quantified from total RNA using strand-specific RT-qPCR and normalized on the level of the U3B snoRNA. Data are presented as mean +/- standard error of the mean (SEM), calculated from three biological replicates. **D.** Strand-specific RT-qPCR analysis of *ctt1* mRNA and *XUT0794* induction in WT cells upon H_2_O_2_ treatment, *ctt1*. mRNA (blue) and *XUT0794* (red) were quantified as above. Relative level of each transcript in the non-induced condition (T0) was set to 1. Mean and SEM values were calculated from three biological replicates.

*ctt1* encodes a catalase, an enzyme required for survival to oxidative stress upon exposure to H_2_O_2_ [33], and it is strongly induced in this condition [34]. Interestingly, we observed that *exo2Δ* cells displayed a slight sensitivity to H_2_O_2_ in addition to the slow growth and temperature sensitivity (S2B-C Figs). This suggests that *ctt1* expression in *exo2Δ* cells might also be impaired upon oxidative stress.

We therefore analyzed *ctt1* mRNA induction in WT and *exo2Δ* cells upon oxidative stress. Northern-blot (Fig 2B) and RT-qPCR kinetics analyses (Fig 2C) showed that *exo2Δ* exhibits a 3-fold reduction in induction rate, with a peak of induction reached 15 minutes after H_2_O_2_ addition vs 10 for the WT (Fig 2C).

Strikingly, we observed that *XUT0794* is also activated upon oxidative stress in a WT context (Fig 2D). Furthermore, its peak of induction is reached very rapidly (5 min), before the *ctt1* mRNA peak (10 min), suggesting that it might be part of a natural attenuation mechanism (feedback loop) for *ctt1* expression.

In summary, our data suggest that *ctt1* induction requires Exo2 activity for maintaining a low level of *XUT0794*, antisense to *ctt1*. Upon induction, the XUT is activated and could modulate expression of *ctt1*, in a similar way as shown for asXUT-associated genes in *S. cerevisiae*, such as *GAL1-10* [19,20].

### *ctt1* attenuation level correlates with antisense *XUT0794* abundance

We designed several experiments in order to test whether *ctt1* attenuation directly depends on antisense *XUT0794*.

Firstly, we overexpressed it in *cis*, in WT cells, using a regulatable *P41nmt1* promoter (S3A Fig). When the promoter is active, *XUT0794* accumulates and *ctt1* is not induced in response to H_2_O_2_ addition (S3A Fig). This demonstrates a causal role of antisense *XUT0794* expression in attenuating *ctt1*. However, in this particular context where *XUT0794* expression is driven by the strong *P41nmt1* promoter, it is difficult to draw up any conclusion about a possible role of the lncRNA itself, as the *ctt1* silencing observed here probably mainly results from transcriptional interference. Note that *P41nmt1-*driven expression of *XUT0794* in *trans*, from a plasmid, failed to attenuate *ctt1* (S3B Fig).

Secondly, we disrupted the *XUT0794* promoter in WT cells using the *ura4* gene (Fig 3A), which is controlled by a promoter much weaker than *P41nmt1*. Surprisingly, *ura4* insertion did not abolish *XUT0794* expression (Fig 3B). However, it resulted in *XUT0794* levels similar to those of the *exo2A* mutant (Fig 3B). Notably, *ctt1* was significantly attenuated in the *ura4-XUT0794* strain, and *ctt1* mRNA levels were similar to those of *exo2Δ* cells (Fig 3C). Hence, there is a positive correlation between *ctt1* attenuation and antisense *XUT0794* levels.

**Fig 3.**
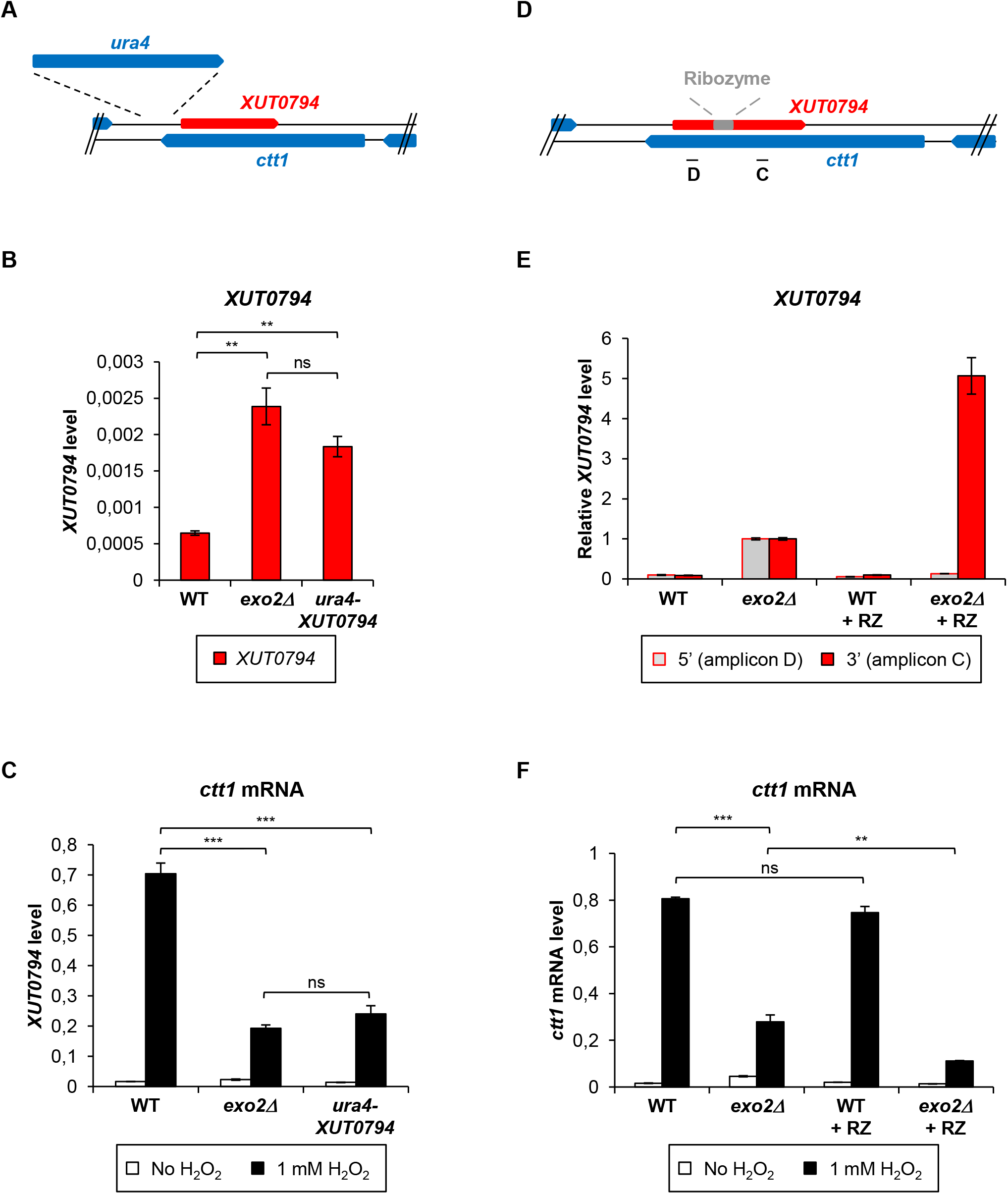
*ctt1* attenuation level correlates with *XUT0794* abundance. **A.** Schematic representation of *ura4* insertion upstream from *XUT0794*. **B.** Analysis of *XUT0794* level upon *ura4* insertion. Strains YAM2400 (WT), YAM2402 (*exo2Δ*) and YAM2817 (*ura4-XUT0794*) cells were grown to mid-log phase in rich YES medium. *XUT0794* level was determined by strand-specific RT-qPCR from total RNA as described in Fig 2C. Data are presented as mean +/− SEM, calculated from three biological replicates. ** *p* < 0.01; ns, not significant upon t-test. **C.** Analysis of *ctt1* mRNA level upon *ura4* insertion. Strains YAM2400 (WT), YAM2402 (*exo2Δ*) and YAM2817 (*ura4-XUT0794*) cells grown to mid-log phase in rich medium were collected before or after addition of 1 mM H_2_O_2_ for 15 min. *ctt1* mRNA level was determined by strand-specific RT-qPCR as described in Fig 2C. Mean and SEM are calculated from three biological replicates. *** *p* < 0.001; ns, not significant upon t-test. **D.** Schematic representation of hammerhead ribozyme (RZ) inserted within *XUT0794*. Self-cleaving RZ was inserted at position 254 of *XUT0794*. Position of qPCR amplicons C and D is indicated. **E.** Analysis of *XUT0794* upstream from and downstream to RZ insertion site. YAM2400 (WT), YAM2402 (*exo2Δ*), YAM2565 (WT + RZ) and YAM2567 (*exo2Δ* + RZ) were grown as described in Fig 3B. Strand-specific RT on *XUT0794* was performed from total RNA using oligonucleotide AMO2069. Oligonucleotides AMO2535-6 (amplicon D) and AMO2069-70 (amplicon C) were used for qPCR detection of *XUT0794* 5′ and 3′ fragment, respectively. Data were normalized on U3B snoRNA. For each amplicon, the normalized level of*XUT0794* in *exo2Δ* was then set to 1. Results are presented as mean +/− SEM, calculated from four biological replicates. **F.** Analysis of *ctt1* mRNA levels. Strains and cultures were in Fig 3C; strand-specific RT on *ctt1* mRNA was performed using oligonucleotide AMO2535; qPCR was performed using AMO2535-6 (amplicon D). Data were normalized on U3B levels and are presented as mean +/− SEM, calculated from four biological replicates. \*\**p*<0.01; \*\*\**p*<0.001; ns, not significant upon t-test.

In a third experiment, we inserted a self-cleaving hammerhead ribozyme (RZ) at position 254/815 of *XUT0794* (Fig 3D) and integrated the construct at the *ctt1* locus in WT and *exo2Δ* strains, without any manipulation of *XUT0794* promoter. In the WT + RZ context, neither the 5′ nor the 3′ fragment of *XUT0794* accumulated (Fig 3E), and *ctt1* induction was similar to WT cells (Fig 3F). In the *exo2Δ* + * RZ context, the 5′ fragment was not detected, but the 3′ fragment accumulated 5x more than in *exo2Δ* without RZ (Fig 3E). This imbalance between the two RNA parts indicates that RZ was efficiently cleaved, the 5′ fragment being presumably degraded [35] while the 3′ fragment accumulated. Importantly, the higher abundance of the 255-815 fragment of *XUT0794* in *exo2Δ* + RZ cells compared to *exo2Δ* without RZ correlated with a significantly stronger attenuation of ctt1 (Fig 3F).

We conclude that the 255-815 fragment of *XUT0794* is sufficient to attenuate *ctt1* and that the level of the attenuation depends on the abundance of the asXUT, which is consistent with the hypothesis that the regulation is mediated by the asXUT but is not an indirect effect of Exo2 inactivation.

### Transcriptional attenuation of *ctt1* in *exo2Δ* cells is characterized by partial RNAPII Ser5 phosphorylation

To determine whether the attenuation of *ctt1* induction occurs at the transcriptional level, we performed RNAPII ChIP experiments in WT and *exo2Δ* cells. Upon oxidative stress, RNAPII occupancy in the mutant showed a significant 2-to 4-fold decrease along the *ctt1* locus (Fig 4A-B), indicating that the attenuation is transcriptional.

**Fig 4.**
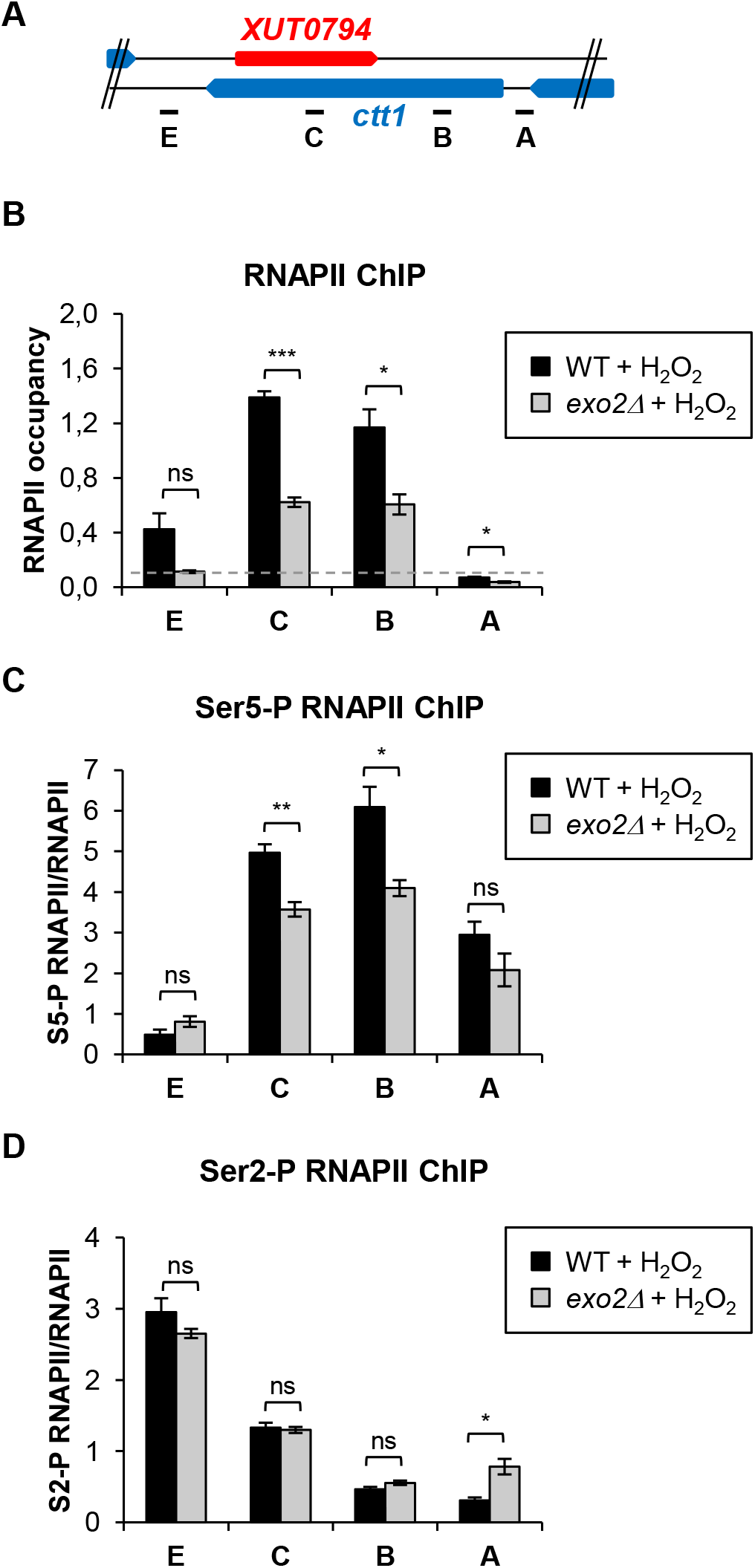
asXUT-mediated attenuation of *ctt1* is transcriptional. **A.** Schematic map of the *ctt1* locus, with positions of the qPCR oligonucleotide pairs. **B.** ChIP analysis of RNAPII occupancy across *ctt1*. Strains YAM2400 (WT) and YAM2402 (*exo2Δ*) were grown as in Fig 2B. After cross-linking, chromatin extraction and sonication, RNAPII was immunoprecipitated using antibody against the CTD of its largest subunit Rpb1. Co-precipitated DNA was purified and quantified by qPCR. Data were normalized on the *act1* signal, which is not controlled to an asXUT (see S4 Fig). The dashed line indicates the background signal observed on an intergenic region of chromosome I used as negative control. Data are presented as mean +/− SEM, from three biological replicates. \**p*<0.05; \*\*\**p*<0.001; ns, not significant upon t-test. **C-D.** ChIP analysis of Ser5-P and Ser2-P occupancy along *ctt1*. RNAPII was immunoprecipitated from the same chromatin extracts as above, using antibody against the Ser5-P **(C)** or the Ser2-P **(D)** form of Rpb1 CTD. Data normalization was as above. For each position of *ctt1*, the ratio between Ser5-P or Ser2-P and total RNAPII is shown. Mean and SEM values were calculated from three biological replicates. \**p*<0.05; \*\**p*<0.01 ns, not significant upon t-test.

Analysis of the distribution of differentially phosphorylated forms of the C-terminal domain (CTD) of Rpb1, the largest subunit of RNAPII, provided further insights into the mechanism of transcriptional attenuation. Ser5-P RNAPII is associated to the early stages of the transcription cycle and predominates in the promoter-proximal region of the gene, while Ser2-P RNAPII is associated to transcription elongation and increases along the gene core [36]. Upon oxidative stress, we observed a 30% decrease of Ser5-P RNAPII in the 5′ and core regions of *ctt1*, in the *exo2Δ* mutant (Fig 4C). In contrast Ser2-P RNAPII occupancy was unaffected (Fig 4D).

Interestingly, we noted that Ser5-P RNAPII levels remain high across the *ctt1* gene body, especially in the *XUT0794* overlapping region (Fig 4C, probe C). This could possibly reflect re-initiation events following collision between convergent RNAPII, in keeping with that *XUT0794* expression is also activated upon oxidative stress (Fig 2D).

In summary, stabilization of *XUT0794* impairs *ctt1* transcription, with less RNAPII loaded on the gene in response to oxidative stress and an additional reduction of Ser5P.

### Attenuation of *ctt1* induction depends on histone deacetylation

Several studies in budding yeast have pointed out the role of HDAC, including the class II HDAC Hda1, in aslncRNA-mediated gene silencing [16,18,19]. To test whether the attenuation of *ctt1* induction involves an HDAC activity, WT and *exo2Δ* cells were treated with trichostatin A (TSA), an inhibitor of class I-II HDAC. When exposed to oxidative stress, TSA-treated *exo2Δ* cells accumulated *ctt1* mRNA to the same level as the control (DMSO-treated) WT strain (Fig 5A). We also noted that the basal levels of *ctt1* mRNA were increased in the TSA-treated WT and *exo2Δ* cells, indicating that *ctt1* repression requires an HDAC activity. Furthermore, both *ctt1* mRNA and *XUT0794* levels in TSA-treated WT cells showed a 2-fold increase compared to the DMSO-treated control after H_2_O_2_ addition (Figs 5A-B). This indicates that the *XUT0794*-associated feedback loop that could modulate *ctt1* expression in WT cells upon exposure to H_2_O_2_ is impaired when HDAC activity is inhibited.

**Fig 5.**
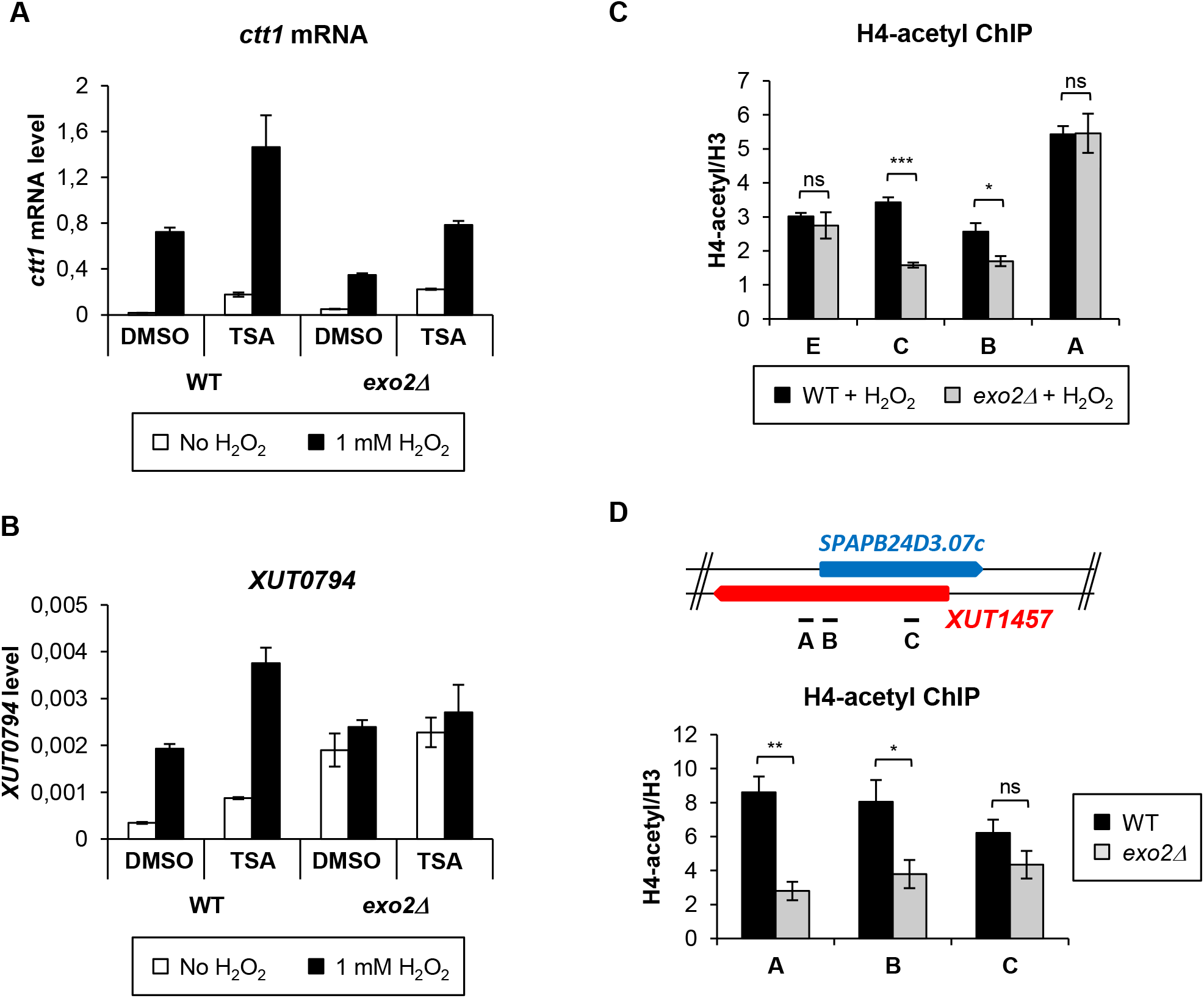
asXUT-mediated attenuation of *ctt1* depends on HDAC. **A-B.** Strand-specific RT-qPCR analysis of *ctt1* mRNA and *XUT0794* levels in the presence of HDAC inhibitor. Strains YAM2400 (WT) and YAM2402 (*exo2Δ*) were grown in rich medium to mid-log phase before addition of 40 μg/ml TSA or equivalent volume of DMSO for 2 hours. TSA-treated and control cells were then collected prior or after addition of H_2_O_2_ for 15 min. *ctt1* mRNA **(A)** and *XUT0794* **(B)** were quantified as described above. Data are presented as mean +/-SEM, calculated from three biological replicates. **C.** ChIP analysis of H4K5/8/12/16 acetylation along *ctt1*. Culture, cross-linking and chromatin extraction were as described in Fig 4B. For each position, data were first normalized on *act1*, then on the level of histone H3, immunoprecipitated from the same chromatin. Data are presented as above. \**p*<0.05; \*\*\**p*<0.001;ns, not significant upon t-test. **D.** ChIP analysis of H4K5/8/12/16 acetylation along the *SPAPB24D3.07c* gene (class 1) in WT and *exo2Δ* cells. YAM2400 (WT) and YAM2402 (*exo2Δ*) cells were grown to mid-log phase in rich medium. Data normalization and presentation was as above. Cross-linking, chromatin extraction, data analysis and presentation were as described above. \**p*<0.05; \*\**p*<0.01; ns, not significant upon t-test.

On the basis of this observation, we predicted histone acetylation along *ctt1* to decrease upon *XUT0794* stabilization. ChIP experiments in *ctt1* induction conditions revealed a significant 50% and 30% reduction of histone H4K5/8/12/16 acetylation and H3K14 acetylation, respectively, in the *exo2Δ* mutant, in the region where ctt1 gene and *XUT0794* overlap (Fig 5C and S5A Fig; probe C). These data support the idea that asXUT-mediated gene attenuation depends on HDAC, resulting in reduced levels of histone acetylation. Importantly, significant histone deacetylation in *exo2Δ* was also detected across *SPAPB24D3.07c*, another class 1 gene (Fig 5D), but not across the class 2 genes *ptb1* and *cuf1* (S5B-C Figs). This indicates that histone deacetylation is not a general feature of all the genes that are transcriptionally down-regulated in *exo2Δ* cells (classes 1-2) but is specific to those with asXUT (class 1).

In an attempt to identify the HDAC involved, we tested the effect of Clr3 (the ortholog of Hda1), Hos2 (a class I HDAC) and Clr6 (the ortholog of class I HDAC Rpd3). As Clr6 exists in at least two distinct complexes (Clr6-CI and -CII), we also tested a specific subunit for each them, namely the ING family protein Png2 (Clr6-CI) and the Sin3 family protein Pst2 (Clr6-CII), respectively [37]. We used null mutants for Clr3, Hos2, Png2 and Pst2, which are non-essential. For Clr6, which is essential, we used the thermo-sensitive *clr6-1* point mutation [38]. Except for *pst2Δ*, we could successfully combine these mutations with *exo2Δ*. Attenuation of *ctt1* was not suppressed in the *exo2Δ clr3Δ, exo2Δ hos2Δ, exo2Δ png2Δ and exo2Δ clr6-1* mutants (S6A-D Figs), indicating that none of the four tested factor is involved in the attenuation mechanism. In contrast, the *png2Δ, pst2Δ* and *clr6-1* single mutants exhibited a strong defect of *ctt1* induction (S6C-E Figs). In addition, the *png2Δ* and *clr6-1* mutations were synergic with *exo2Δ* (S6C-D Figs). This indicates that Exo2 and the Clr6 HDAC complexes are required for efficient *ctt1* induction, but act independently.

In conclusion, XUT-mediated attenuation of *ctt1* requires a HDAC activity, suggesting that mechanisms of regulation of gene expression by lncRNAs have been conserved across the yeast clade.

### Role of Set2-dependent H3K36me3 in *ctt1* attenuation

During transcription, elongating RNAPII recruits the histone methyltransferase (HMT) Set2, which methylates H3K36 across the gene body [39,40]. Set2-mediated H3K36 trimethylation (me3) then promotes HDAC recruitment and histone deacetylation [41,42,43], in order to suppress spurious intragenic transcription initiation [44,45].

The observation of decreased histone acetylation levels across the *ctt1* gene body in *exo2Δ* cells (Fig 5C) prompted us to test the role of Set2 and H3K36 methylation in the regulation. In WT cells, upon oxidative stress, we observed a peak of Set2 occupancy and H3K36me3 in the region overlapping *XUT0794* (probe C; Figs 6A-B), confirming that Set2 is recruited when *ctt1* expression is induced. In the *exo2Δ* mutant, H3K36me3 levels were significantly increased, in some positions of *ctt1*, including the 5′ region (probe B) and the 3′ extremity (probe E). Surprisingly, in the region overlapping *XUT0794* (probe C), where Set2 occupancy and histone deacetylation are the most pronounced (Figs 5C and 6A), the difference with the WT control was not statistically significant (Fig 6B). Perhaps a local change of H3K36me3 does not result into histone deacetylation at that position but to the nearby region.

**Fig 6.**
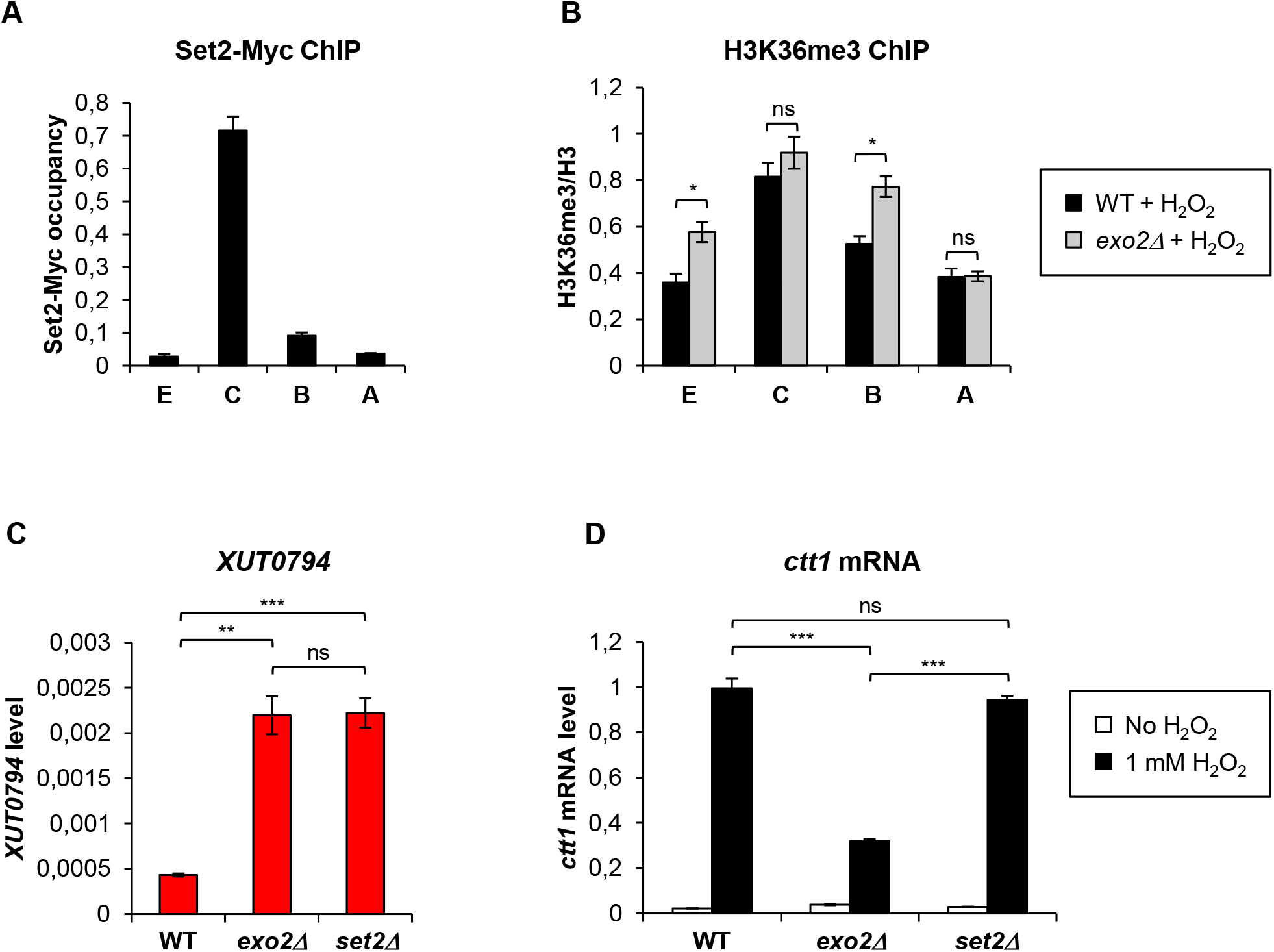
Role of Set2-dependent H3K36me3 in *ctt1* attenuation. **A.** Set2 is recruited to *ctt1* upon oxidative stress. YAM2816 (Set2-13Myc [60]) cells were grown as in Fig 2B. Cross-linking and chromatin extraction were as described in Fig 4. Data are presented as mean +/-SEM, calculated from three biological replicates. **B.** ChIP analysis of H3K36 trimethylation (me3) along *ctt1* in *exo2Δ* cells. Strains YAM2400 (WT) a698 nd YAM2402 (*exo2Δ*) were grown as in Fig 2B. Cross-linking and chromatin extraction were as described in Fig 4. Data analysis was performed as described in Fig 5C. Average values and SEM were calculated from three biological replicates. \**p*<0.05; ns, not significant upon t-test. **C.** Analysis of *XUT0794* level in *set2Δ* cells. Strains YAM2400 (WT), YAM2402 (*exo2Δ*) and YAM2797 (*set2Δ*) were grown to mid-log phase in rich medium. XUT0794 level was quantified from total RNA as described in Fig 2C. Data are presented as mean +/− SEM, calculated from three biological replicates. ** *p* < 0.01; \*\*\**p* < 0.001; ns, not significant upon t-test. **D.** Analysis of *ctt1* mRNA induction in *set2Δ* cells. Strains YAM2400 (WT), YAM2402 (*exo2Δ*) and YAM2797 (*set2Δ*) were grown as in Fig 2B. ctt1 mRNA was quantified by strand-specific RT-qPCR as described in Fig 2C. Data are presented as above. *** *p* < 0.001; ns, not significant upon t-test.

In parallel, we analyzed the effect of Set2 inactivation on *XUT0794* expression and *ctt1* mRNA induction. We could only characterize single mutants, as despite our efforts, we failed to combine *set2Δ* and *exo2Δ*, suggesting that the double mutant is lethal. Strikingly, we found that *set2Δ* cells accumulate *XUT0794* into levels similar to the *exo2Δ* mutant (Fig 6C). However, ctt1 induction was found to be normal in the *set2Δ* context (Fig 6D). These data indicate that the *XUT0794*-associated regulation of *ctt1* is impaired when Set2 is inactivated.

In summary, Set2 is recruited to *ctt1* upon oxidative stress. In absence of Set2, *XUT0794* accumulation and *ctt1* attenuation are decoupled, indicating that Set2 is required for the *XUT0794-* associated regulation of *ctt1*.

### Dcrl regulates *ctt1* induction independently of Exo2

The data presented above show that the Exo2-dependent RNA decay controls *ctt1* induction and restricts the level of the antisense *XUT0794*. We asked whether other RNA processing factors could be involved in the regulation. We tested the role of Dicer (Dcr1), involved in RNAi.

As shown in Fig 7A, ctt1 attenuation is not suppressed in the *exo2Δ dcr1Δ* double mutant. Rather, ctt1 was found to be attenuated in the *dcr1Δ* single mutant, and we observed a synergic effect in the *exo2Δ dcr1Δ* double mutant. On the other hand, Dicer overexpression in *exo2Δ* cells had no impact on *ctt1* attenuation (S7A-C Figs). These data indicate that Exo2 and Dcr1 control *ctt1* induction through distinct mechanisms. This is further supported by the observation that *XUT0794* levels are unchanged in *exo2Δ dcr1Δ* cells compared to *exo2Δ* cells (Fig 7B) and by ChIP experiments showing that in oxidative stress conditions, RNAPII (Fig 7C) and H3K36me3 (S7D Fig) levels are normal in *dcr1Δ* cells (Fig 7C) while histone H4K5/8/12/16 acetylation is strongly reduced across the whole *ctt1* locus, including the promoter region (Fig 7D).

**Fig 7.**
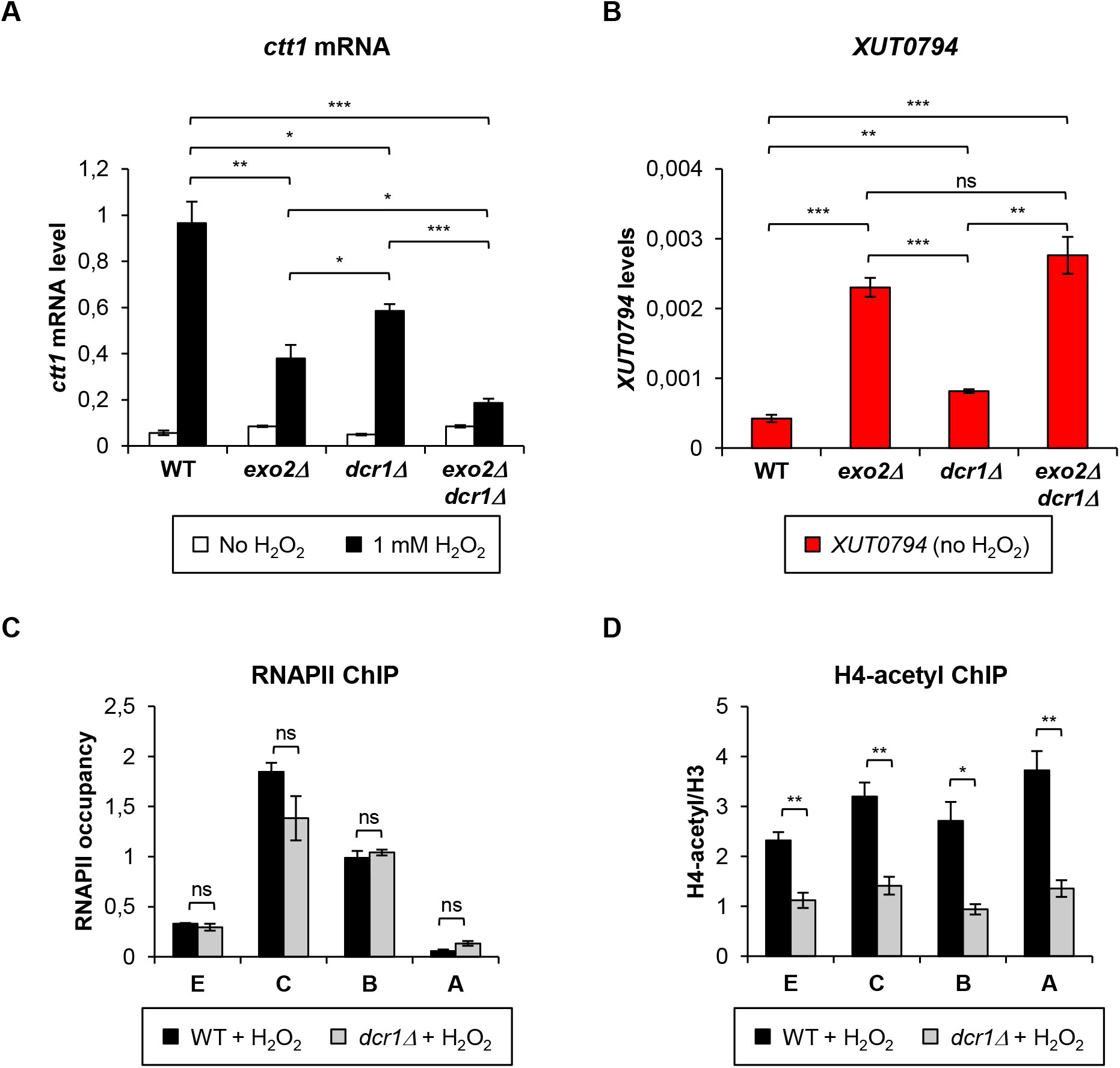
Dicer controls *ctt1* induction independently of Exo2. **A.** Analysis of *ctt1* mRNA induction in WT, *exo2Δ, Dcr1Δ* and *exo2Δ Dcr1Δ* cells. Strains YAM2400 (WT), YAM2402 (*exo2Δ*), YAM2406 (*Dcr1Δ*) and YAM2404 (*exo2Δ Dcr1Δ*) were grown as described in Fig 2B. *ctt1* mRNA was quantified by strand-specific RT-qPCR as described in Fig 2C. Data are presented as mean +/− SEM, calculated from three biological replicates. \**p* < 0.05; \*\**p* < 0.01; \*\*\**p* < 0.001 upon t-test. **B.** Analysis of *XUT0794* level in in WT, *exo2Δ, Dcr1Δ* and *exo2Δ Dcr1Δ* cells. Strains YAM2400 (WT), YAM2402 (*exo2Δ*), YAM2406 (*Dcr1Δ*) and YAM2404 (*exo2Δ Dcr1Δ*) were grown to mid-log phase in rich medium. *XUT0794* was quantified by strand-specific RT-qPCR as described in Fig 2C. Data are presented as mean +/− SEM, calculated from three biological replicates. \*\**p* < 0.01; \*\*\**p* < 0.001; ns, not significant upon t-test. **C.** ChIP analysis of RNAPII occupancy across *ctt1* in *dcr1Δ* cells. Strains YAM2400 (WT) and YAM2406 (*dcr1Δ*) were grown as in Fig 2B. Cross-linking, chromatin extraction and data analysis were as described in Fig 4B. Data are presented as mean +/− SEM, calculated from three biological replicates, ns, not significant upon t-test. **D.** ChIP analysis of H4K5/8/12/16 acetylation (H4-acetyl) along *ctt1* in *dcr1Δ* cells. Strains YAM2400 (WT) and YAM2406 (*dcr1Δ*) were grown as in Fig 2B. Cross-linking and chromatin extraction were as described in Fig 4B. Data analysis was performed as described in Fig 5C. Data are presented as mean +/− SEM, calculated from three biological replicates. \**p*<0.05; \*\**p*<0.01 upon t-test.

Thus, Exo2 and Dcr1 regulate *ctt1* induction through independent mechanisms, which is consistent with the observation that asXUTs are globally not targeted by RNAi in *S. pombe* [30].

## DISCUSSION

Previous studies in different eukaryotic models have shown that aslncRNAs can regulate sense gene expression [14]. However, the molecular bases for aslncRNAs-mediated regulation remain largely unknown. In budding and fission yeasts, aslncRNAs are actively degraded by the Xrn1/Exo2-dependent cytoplasmic 5′-3′ RNA decay pathway [28,29,30]. These Xrn1/Exo2-sensitive aslncRNAs are named XUTs [28]. In budding yeast, asXUTs stabilization was shown to result into transcriptional attenuation of a subset of genes, referred to as class 1 [28]. Whether such an asXUT-associated regulation is conserved in other organisms was unknown.

Here, we used NET-Seq to identify genes showing transcriptional attenuation upon stabilization of their paired-asXUT (class 1) in *S. pombe*. Importantly, asXUT presence and sense gene attenuation in *exo2Δ* are not independent, supporting the idea that the regulation is mediated by the stabilized asXUTs and is not a side effect of Exo2 inactivation. However, additional mechanistic analyses are required to confirm this hypothesis.

In a previous study, we reported that the asXUT-associated genes are globally less transcribed than the ‘solo’ ones (without asXUT), displaying an hypoacetylated promoter and hyperacetylation across the gene body [30]. Here we show that the asXUT-associated genes can be separated in two distinct subgroups, namely class 1 (attenuated upon asXUT stabilization in *exo2Δ*) and class 3 (unchanged in *exo2Δ*). Class 1 corresponds to highly transcribed genes showing prominent nucleosome depletion and high histone acetylation levels at the promoter. In addition, class 1 displays high TSS-proximal antisense transcription, suggesting that the TSS region could be a possible determinant for aslncRNA-mediated regulation. In contrast, class 3 is weakly transcribed, and displays poor promoter-proximal nucleosome depletion and low histone acetylation. Upon stabilization of asXUTs, transcription of class 1 drops down to the basal levels of class 3. This suggests the existence of a regulatory threshold, *ie* asXUTs would modulate expression of their associated sense genes, only if expression is above this threshold. This hypothesis contrasts with a previous model based on the analysis of sense-antisense RNA levels in budding yeast, which proposed that antisense-mediated repression would be restricted to low sense expression [46].

Our data suggests that a subset of asXUTs could regulate gene expression at the transcriptional level, reducing sense transcription, as previously shown in *S. cerevisiae* [18,28]. XUTs could also act at other steps of the gene expression process, especially at the post-transcriptional level. In this regard, aslncRNAs have been shown to modulate protein production in response to osmotic stress in *S. pombe* [23]. In *S. cerevisiae*, disruption of several aslncRNAs results into increased protein synthesis from their paired-sense mRNAs, indicating a role of these aslncRNAs in the control of protein abundance [47]. Future investigations will be required to explore the regulatory potential of asXUTs and to determine the step(s) of the gene expression process they act on.

To get insights into the mechanism by which asXUTs could attenuate gene expression, we selected a class 1 representative, the catalase-coding gene *ctt1*, for further characterization. Induction of *ctt1* in response to oxidative stress was attenuated in *exo2Δ* cells, correlating with antisense *XUT0794* accumulation. Our data indicate that *ctt1* attenuation in Exo2-deficient cells occurs at the transcriptional level and is mediated by HDAC activity. Our attempts to identify the HDAC involved in the attenuation mechanism were unsuccessful, most likely due to redundancy of HDAC activities, considering the results obtained upon TSA treatment. On the other hand, we show that the Clr6CI-ll complexes (the homolog of Rpd3L and Rpd3S, respectively) are required for *ctt1* induction (S6C-E Fig). The mechanism of HDAC recruitment also remains to be determined. Although we cannot formally exclude a direct recruitment by the asXUT itself, the HDAC is probably recruited through the Set2-dependent H3K36me3 marks. Consistent with this hypothesis, the most hypoacetylated region of *ctt1* corresponds to the peak of Set2 occupancy and H3K36me3. Furthermore, at the RNA level, the loss of HDAC activity and the inactivation of Set2 have a similar effect, decoupling *XUT0794* accumulation and *ctt1* regulation.

Whether *ctt1* attenuation in *exo2Δ* cells depends on the stabilized asXUT per se and/or on the act of antisense transcription remains unsolved to date. On one hand, the ribozyme experiment shows that *ctt1* attenuation level positively correlates with *XUT0794* abundance (Figs 3E-F), in a context where the promoter of the XUT (and presumably the level of antisense nascent transcription) remains unchanged, which is consistent with a regulation mediated by the RNA. On the other hand, the local increase of H3K36me3 across the *ctt1* gene body in *exo2Δ* cells (Fig 6B) suggests that *ctt1* regulation could also depend on transcriptional interference. In fission yeast, gene repression by transcriptional interference requires Set2 and the Clr6CII complex [48]. Here, we show that Set2 and Clr6CII have different effects on *ctt1* induction: it is normal in Set2-deficient cells (Fig 6D) but attenuated upon inactivation of Clr6CII components (S6E Fig). This suggests that the roles of Set2 and Clr6CII might differ from a gene to another.

The model of a RNA-mediated regulation raises a key mechanistic question: how could an asXUT, which in all likelihood accumulates in the cytoplasm in *exo2Δ* cells, act in the nucleus to regulate the transcription of their paired-sense genes? This remains unknown to date. One possibility is that the stabilized XUTs could shuttle between the cytoplasm to the nucleus, as shown for tRNAs in budding yeast [49,50] and also in mammalian cells [51].

Most of the effects we described have been observed in mutant cells. But does antisense *XUT0794* play any role in WT cells? Interestingly, we observed that *XUT0794* is also rapidly induced in WT cells after H_2_O_2_ addition, suggesting that it could participate in the modulation of *ctt1* induction. In this respect, a recent study in human fibroblasts identified a class of stress-induced aslncRNAs, which are activated upon oxidative stress [8], suggesting that aslncRNAs induction might be part of a conserved response to oxidative stress in Eukaryotes. Demonstrating that *XUT0794* plays a direct role in the modulation of *ctt1* during oxidative stress relies, among others, on the ability to block its expression. Unfortunately, none of the strategies we used in this work succeeded into blocking *XUT0794*. Additional work will be required to implement in fission yeast other techniques developed in *S. cerevisiae* to strand-specifically block aslncRNA synthesis [47,52], which remains technically challenging yet. For instance, at some loci, the CRISPR interference approach is not strand-specific and results in the production of novel isoforms of the targeted aslncRNA [53].

Efficient Induction of *ctt1* upon oxidative stress depends on multiple factors [54]. In addition to Exo2, we showed that Dcr1 also contributes to *ctt1* induction. Our data indicate that Dcr1 and Exo2 regulate *ctt1* through distinct mechanisms, which is consistent with the observations that asXUTs are not targeted by Dicer in fission yeast [30]. Interestingly, Dcr1 was recently shown to promote efficient termination of a set of highly transcribed genes, corresponding to sites of replication stress and DNA damage [55], and *ctt1* belongs to this set of Dcr1-terminated genes. Thus, one possibility could be that Dcr1 regulates *ctt1* induction at the level of transcription termination. However, our ChIP data did not reveal any significant change of RNAPII occupancy at the 3′ extremity of *ctt1* in *dcr1Δ* cells, upon oxidative stress (Fig 7C). Additional analyses are therefore required to decipher the mechanism by which Dcr1 regulates *ctt1* induction.

In conclusion, our work in budding and fission yeasts shows that the cytoplasmic 5′-end RNA decay plays a key role in controlling aslncRNAs endowed with regulatory potential. Given the high conservation of Xrn1 in Eukaryotes, it is tempting to speculate that asXUTs and their regulatory activity are conserved in higher eukaryotes, contributing in buffering genome expression, and adding another layer to the complexity of gene regulation.

## MATERIALS & METHODS

### Yeast strains, plasmids and media

All the strains used in this study are listed in S5 Table. Mutant strains were constructed by meiotic cross or transformation, and verified by PCR on genomic DNA and/or RT-qPCR. Plasmid pAM353 for expression of *XUT0794* in *trans* was constructed by cloning *XUT0794* in the Sail site of pREP41 (*ars1 LEU2 P41nmt1*). Sanger sequencing confirmed the correct orientation of the insert and the absence of mutation. Hammerhead ribozyme [56] was inserted in *XUT0794* by two-step PCR, giving a 3.2 Kb final product corresponding to *ctt1* mRNA coordinates +/− 500 bp that was cloned in pREP41. After verification of absence of additional mutations by Sanger sequencing, the ribozyme-containing construct was excised and transformed in the YAM2534 strain (*ctt1::ura4*). Transformants were selected on 5-FOA plates and analyzed by PCR on genomic DNA. Deletion of *exo2* was performed subsequently.

Strains were grown at 32°C to mid-log phase (OD_595_ 0,5) in YES or EMM-L medium. For *ctt1* induction, 1 mM H2O2 was added for 15 minutes [34], or different time points for analysis of kinetics of induction. Expression from *P41nmt1* was repressed by growing cells in EMM-L + 15 μM thiamine for 24 hours.

### NET-Seq

NET-Seq libraries were constructed from biological duplicates of YAM2507 (*exo2Δ rpb3-flag*) cells and sequenced as previously described [30]. Libraries for the WT strain YAM2492 (*rpb3-flag*) were described in the same previous report [30].

After removal of the 5′-adapter sequence, reads were uniquely mapped to the reference genome (ASM294v2.30) using version 0.12.8 of Bowtie [57], with a tolerance of 2 mismatches.

Differential analysis was performed between the IP samples from WT and *exo2Δ* using DESeq [58]. Genes showing significant decrease (*P*-value <0.05, adjusted for multiple testing with the Benjamini-Hochberg procedure) in the mutant were defined as class 1 & 2.

Raw sequences have been deposited to the NCBI Gene Expression Omnibus (accession number GEO: GSE106649). A genome browser for visualization of NET-Seq processed data is accessible at http://vm-gb.curie.fr/mw3.

### Total RNA extraction

Total RNA was extracted from exponentially growing cells using standard hot phenol procedure, resuspended in nuclease-free H_2_O (Ambion) and quantified using a NanoDrop 2000c spectrophotometer.

### Northern blot

10 μg of total RNA were loaded on denaturing 1.2% agarose gel and transferred to Hybond^™^-XL nylon membrane (GE Healthcare), *ctt1* mRNA and U3B were detected using AM02063 and AM02081 oligonucleotides, respectively (see S6 Table). ^32^P-labelled probes were hybridized overnight at 42°C in ULTRAhyb®-Oligo hybridization buffer (Ambion). Quantitation used a Typhoon Trio Phosphorlmager and the ImageQuant TL v5.2 sofware (GE Healthcare).

### Strand-specific RT-qPCR

Strand-specific reverse transcription (RT) reactions were performed from at least three biological replicates, using 1 μg of total RNA and the SuperScript^®^II Reverse Transcriptase kit (Invitrogen), in the presence of 6,25 μg/ml actinomycin D. For each sample, a control without RT was included. Subsequent quantitative real-time PCR were performed on technical duplicates, using a LightCycler^®^ 480 instrument (Roche). Oligonucleotides used are listed in S6 Table.

### ChIP

ChIP analysis was performed from three biological replicates, for each strain. Exponentially growing (OD_595_ 0,5) cells were fixed for 10 minutes at room temperature using formaldehyde (1% final concentration), then glycine was added (0,4 M final concentration) for 5 minutes. Chromatin was sonicated using a Bioruptor^®^ sonication device (Diagenode). Antibodies used were 8WG16 (Covance) for RNAPII, H14 (Covance) for RNAPII S5-Pho, 3E10 (Millipore) for RNAPII S2-Pho, ab1791 (Abcam) for histone H3, 05-1355 (Millipore) for acetyl-H4 (Lys5/8/12/16), 07-353 (Millipore) for acetyl-H3 (Lys14), ab9050 (Abcam) for H3K36me3 and 9E10 (Protein Expression and Purification Core Facility, Institut Curie) for Myc. Quantitative real-time PCR were performed in technical duplicates on a StepOnePlus^™^ machine (Applied Biosystems) or on a LightCycler^®^ 480 instrument (Roche). Oligonucleotides used are listed in S6 Table.

## ACKNOWLEDGMENTS

We would like to thank Fred Winston for the Set2-Myc strain and Danesh Moazed for the *Dcr1OE* vector. We also thank Thomas Rio Frio, Sylvain Baulande, Patricia Legoix-Né and Virginie Raynal (NGS platform, Institut Curie). We are grateful to Nicolas Vogt and Ugo Szachnowski for assistance. We thank all the members of our labs for discussions and critical reading of the manuscript.

## SUPPORTING INFORMATION LEGENDS

**S1 Fig. Antisense XUT stabilization induces transcriptional attenuation of a class of highly expressed genes. A.** Global RNAPII transcription in WT and *exo2Δ* cells. Density plot of *exo2Δ*/WT NET-Seq signal ratio for mRNAs (blue), sn(o)RNAs (black) and XUTs (red). **B.** Metagene view of NET-Seq signals along class 1-4 genes in WT cells. For each class, normalized signal (tag/nt, log_2_) along mRNA transcription start site (TSS) +/− 1000 nt (sense strand) and the antisense (as) strand were piled up, in a strand-specific manner. Average signal for each strand was plotted for class 1 (red), 2 (blue), 3 (green) and 4 (black). The shading surrounding each line denotes the 95% confidence interval. **C.** Box-plot of NET-Seq signal (tag/nt, log_2_) for XUTs antisense to class 1 and class 3 genes in WT and *exo2Δ* (*Δ*) cells. **D.** Metagene view of nascent antisense transcription (NET-Seq) signal around the sense gene TSS of class 1-4 genes, in WT cells. The shading surrounding each line denotes the 95% confidence interval. **E** Metagene of H3 levels for class 1-4 genes in WT cells. The analysis was performed using previously published ChIP-Seq data [30]. Metagene representation of signal for class 1 (red), class 2 (blue), class 3 (green) and class 4 (black) was performed as above, in a strand-unspecific manner. The shading surrounding each line denotes the 95% confidence interval. **F.** Metagene view of H3K14 acetylation for class 1-4 genes in WT cells. ChIP-Seq libraries construction and sequencing were previously described [30]. Metagene representation of signal for each class of genes was performed as above, using ratio of coverage (log_2_) for H3K14ac and H3.

**S2 Fig. Stabilization of *XUT0794* antisense to *ctt1* correlates with H_2_O_2_ sensitivity in *exo2Δcells*. A.** Snapshot of total RNA-Seq signals along the *ctt1* locus in WT and *exo2Δ* cells. RNA-Seq data were previously published [30]. Densities (tag/nt) for the + and - strands are visualized in the upper and lower panels, respectively. The signals for the WT and the *exo2Δ* strains are represented as black and grey lines, respectively. **B.** Exo2-deficient cells display a slow growth phenotype. YAM2400 (WT) and YAM2402 (*exo2Δ*) cells were grown in rich (YES) medium, at 32°C. OD_595_ was measured every hour. OD_595_ at time 0 was set to 1, for each strain. Data are expressed in a log scale. **C.** Loss of Exo2 confers sensitivity to hydrogen peroxide. Serial 1:10 dilutions of YAM2400 (WT) and YAM2402 (*exo2Δ*) cells were dropped on solid rich medium (YES) containing or not 2 mM H_2_O_2_. Plates were incubated at the indicated temperature for 3-4 days.

**S3 Fig. Effect of *XUT0794* overexpression in *cis* and in *trans*. A.** Attenuation of *ctt1* mRNA upon overexpression of *XUT0794* in *cis*. Strains YAM2400 (WT) and YAM2474 (*P41nmt1-XUT0794*) were grown for 24 hours to mid-log phase in EMM medium +/− 15 μM thiamine, before addition of H_2_O_2_ for 15 min. Levels of *XUT0794* (red) and *ctt1* mRNA (blue) were quantified from total RNA using strand-specific RT-qPCR and normalized on the level of the U3B snoRNA. Data are presented as mean +/− SEM from three biological replicates. **B.** Induction of *ctt1* mRNA upon *XUT0794* overexpression in *trans*. YAM2475 (empty vector) and YAM2476 (pAM353; *P41nmt1-XUT0794*) cells were grown for 24 hours to mid-log phase in EMM-L medium, before addition of H_2_O_2_ for 15 min. Determination of *XUT0794* and *ctt1* mRNA levels and data presentation are as above.

**S4 Fig. Transcription of *act1* is not controlled by an asXUT**. Snapshot of total (input) and nascent (IP) NET-Seq signals along the *act1* gene in WT (upper panels) and *exo2Δ* (lower panels) cells. In each panel, the signal corresponding to the sense (+) and antisense (−) strand is shown in blue and pink, respectively. Blue arrows and boxes represent the mRNAs and coding sequences, respectively. NET-Seq data for the WT strain were previously described [30]. The snapshot was produced using VING [59].

**S5 Fig. Histone deacetyation in *exo2Acells* is specific of class 1. A.** ChIP analysis of H3K14 acetylation along *ctt1*. Culture, cross-linking and chromatin extraction were as described in Fig 4B. For each position, data were first normalized on *act1*, then on the level of histone H3, immunoprecipitated from the same chromatin. Data are presented as mean +/− SEM, calculated from three biological replicates. \**p*<0.05; \*\**p*<0.01; ns, not significant upon t-test. **B-C**. ChIP analysis of H4 K5/8/12/16 acetylation along the class 2 genes *cuf1* (**B**) and *ptb1* (**C**) in WT and *exo2Δ* cells. Cultures were as described in Fig 5D. Cross-linking, chromatin extraction, data analysis and presentation were as above, ns, not significant upon t-test.

**S6 Fig. Analysis of *ctt1* attenuation in HDAC mutants. A.** Effect of the Clr3 class II HDAC. YAM2400 (WT), YAM2402 (*exo2Δ*), YAM2407 (*clr3Δ*) and YAM2444 (*exo2Δ clr3Δ*) cells were grown in rich medium before (white) or after addition of 1 mM H_2_O_2_ for 15 minutes (black), *ctt1* mRNA levels were quantified by strand-specific RT-qPCR from total RNA as described in Fig 2C. Data are presented as mean +/− SEM from three biological replicates. \**p*<0.05; \*\**p*<0.01; ns, not significant upon t-test. **B.** Effect of Hos2 class I HDAC. Same as above using YAM2400 (WT), YAM2402 (*exo2Δ*), YAM2471 (*hos2Δ*) and YAM2472 (*exo2Δhos2Δ*). \**p*<0.05; \*\*\**p*<0.001; \*\*\**p*<0.001; ns, not significant upon t-test. **C.** Effect of the Png2 subunit of the ClrβCI complex. Same as above using YAM2400 (WT), YAM2402 (*exo2Δ*), YAM2561 (*png2Δ*) and YAM2562 (*exo2Δ png2Δ*). \**p*<0.05; \*\**p*<0.01; ***p<0.001; ns, not significant upon t-test. **D.** Effect of Clr6 class I HDAC. Same as above using YAM2400 (WT), YAM2402 (*exo2Δ*), YAM2798 (clr6-1) and YAM2814 (clr6-1 *exo2Δ*). \**p*<0.05; \*\*\**p*<0.001 upon t-test. **E.** Effect of the Pst2 subunit of the Clr6CII complex. Same as above using YAM2400 (WT), YAM2402 (*exo2Δ*) and YAM2815 (*pst2Δ*). \**p*<0.05; \*\**p*<0.01; \*\*\**p*<0.001 upon t-test.

**S7 Fig. Dicer controls *ctt1* induction independently of Exo2. A-C.** WT cells with pREP-nmt1/LEU2 empty vector (pDM829, vector), and *exo2Δ* cells with pREP-nmt1/LEU2 or pREPNFLAG-Dcr1 (pDM914, Dcr1) plasmids [61] were grown to mid-log phase in EMM-Leu medium, before addition of H_2_O_2_ for 15 min. *ctt1* mRNA (**A**), *XUT0794* (**B**) and *Dcr1* mRNA (**C**) were quantified from total RNA using strand-specific RT-qPCR and normalized on the level of the U3B snoRNA. Average values and SEM were calculated from three biological replicates. \*\**p*<0.01; \*\*\**p*<0.001 ns, not significant upon t-test. **D.** ChIP analysis of H3K36 trimethylation (H3K36me3) along *ctt1* in *dcr1Δ* cells. Strains YAM2400 (WT) and YAM2406 (*dcr1Δ*) were grown as in Fig 2B. Cross-linking and chromatin extraction were as described in Fig 4B. Data analysis was performed as described in Fig 5C. Data are presented as mean +/− SEM, calculated from three biological replicates, ns, not significant upon t-test.

**S1 Table. List of class 1 genes.**

**S2 Table. List of class 2 genes.**

**S3 Table. List of class 3 genes.**

**S4 Table. List of class 4 genes.**

**S5 Table. Yeast strains.**

**S6 Table. Oligonucleotides.**

